# A polyetic modelling framework for plant disease emergence

**DOI:** 10.1101/2020.04.23.057372

**Authors:** Laetitia Willocquet, S. Savary, B.A. McDonald, A. Mikaberidze

## Abstract

Plant disease emergences have dramatically increased recently as a result of global changes, especially with respect to trade, host genetic uniformity, and climate change. A better understanding of the conditions and processes determining epidemic outbreaks caused by the emergence of a new pathogen, or pathogen strain, is needed to develop strategies and inform decisions to manage emerging diseases. A polyetic process-based model is developed to analyse conditions of disease emergence. This model simulates polycyclic epidemics during successive growing seasons, the yield losses they cause, and the pathogen survival between growing seasons. This framework considers an immigrant strain coming into a system where a resident strain is already established. Outcomes are formulated in terms of probability of emergence, time to emergence, and yield loss, resulting from deterministic and stochastic simulations. An analytical solution to determine a threshold for emergence is also derived. Analyses focus on the effects of two fitness parameters on emergence: the relative rate of reproduction (speed of epidemics), and the relative rate of mortality (decay of population between seasons). Analyses revealed that stochasticity is a critical feature of disease emergence. The simulations suggests that: (1) emergence may require a series of independent immigration events before a successful invasion takes place; (2) an explosion in the population size of the new pathogen (or strain) may be preceded by many successive growing seasons of cryptic presence following an immigration event, and; (3) survival between growing seasons is as important as reproduction during the growing season in determining disease emergence.

## 1. INTRODUCTION

The emergence of disease in plant populations has important impacts on both agricultural production and natural ecosystems (Anderson *et al.*, 2004; Lucas, 2017). While emerging plant diseases threaten biodiversity and the entire range of services contributed by plants to the biosphere (Anderson *et al.*, 2004), the emergence of plant diseases constitutes an immediate threat to food security, from local to global scales, because of the losses in production, and also because losses to plant disease affect food access (economic or physical) and the quality of food (Savary *et al.* (2017). The literature provides growing evidence that plant disease emergences have dramatically increased recently, as a result of global changes in trade, host genetic uniformity, and climate (Anderson *et al.*, 2004; Fisher *et al.*, 2012; McDonald and Stukenbrock, 2016; Paini *et al.*, 2016).

A relatively recent example of emergence of new pathogen strains is the introduction into Europe of a strain carrying the A2 mating type of *Phythophthora infestans*, the causal agent of potato late blight (Zwankhuizen and Zadoks, 2002; Lucas, 2017). The emergence of this strain and its lineages, both resistant to metalaxyl and more aggressive, led to more diversified, sexually reproducing, pathogen populations, and increased disease intensity in Europe (Goodwin *et al.*, 1996). Stem rust of wheat is another example. Stem rust epidemics, which were common in the USA during the first half of the last century, became rare after the pathogen (*Puccinia graminis* f. sp. *tritici*) was controlled by combining the deployment of new resistance genes in wheat varieties with the eradication of barberry, which is the alternate host on which the pathogen reproduces sexually (Roelfs, 1978; 1985). In 1998, new races of this pathogen (called Ug99) were detected in Uganda that were virulent against resistance genes present in wheat varieties widely grown in East Africa, leading to local but severe epidemics in the region (Singh *et al.*, 2015). International efforts to generate and deploy resistant varieties helped to limit impacts from races of these new lineages (Singh *et al.*, 2015), but the recent detection of stem rust in different parts of Europe is now threatening wheat production in this part of the world (Saunders *et al.*, 2019). A third and recent example of strain introduction is that of *Puccinia striiformis* f. sp. *tritici*, the causal agent of stripe (yellow) rust of wheat, into North-Western Europe in 2011 (de Vallavieille-Pope *et al.*, 2018) which caused serious epidemics.

An example of emergence of a new pathogen is *Pyricularia graminis-tritici*, the cause of wheat blast. The disease was restricted to South America until 2016, when the pathogen was accidentally introduced and caused a severe outbreak in South Asia (Ceresini *et al.*, 2018). Another example of new pathogen emergence is the Asian soybean rust, caused by *Phakopsora pachyrhizi*, which was introduced into South America at the beginning of this century and has since severely impacted soybean production on that continent (Lucas, 2017). Rhizomania is a virus disease of sugar beet that was first detected in the United Kingdom in 1987 and has since spread, resulting in increasing numbers of epidemics (Gilligan *et al.*, 2007). Other recent examples of disease emergence with very disastrous impacts on perennial crops include huanglongbing on citrus in the New World (Gottwald, 2010) and *Xyllela fastidiosa* on olive trees in Southern Europe (Saponari et al, 2019).

Disease emergence may be associated with changes in the environment, especially, human-made changes. A much-debated example is the case of fusarium head blight of wheat (wheat scab), which has been associated with the maize-wheat rotation, and with no-till practices (Zadoks and Schein, 1979; McMullen *et al.*, 2012). Another example is that of false smut of rice, which has been associated with the cultivation of hybrid rice (Savary *et al.*, 2017). A third example of environmental change-driven emergence is that of *Sclerotium rolfsii*, a tropical pathogen on legumes (among many other hosts) becoming prevalent in the state of New York as a result of warming climate (S. Pethybridge, Personal Communication).

In their seminal article, Heesterbeek and Zadoks (1987) proposed a mathematical theory of pandemics, with three phases: zero-order, first-order, and second-order epidemics. This theory considers two groups of processes, the spatial spread of disease and the accumulation of disease cycles within and across crop cycles, to analyse pandemics. While the zero-order epidemic is field-bound and polycyclic, the first-order epidemic is area-bound and polycyclic, and the second-order is both continental and polyetic. The present article is a response, some thirty years later, to this article. Figure 1 represents a synthesis of processes which may be associated with disease emergence, organised in three paths. Path 1 is the invasion of a new pathogen into an ecosystem, through introduction, establishment, and spread. Path 1 is exemplified by the wheat blast epidemic in Bangladesh. Path 2 is the emergence of disease in response to environmental changes in an ecosystem, where environmental changes lead to disease intensification, further leading to disease spread within entire (agro)systems. Path 2 is illustrated by fusarium head blight of wheat or false smut of rice. Path 3 is the emergence of new strains through evolutionary processes. Path 3 is illustrated by wheat stem rust in Sub-Saharan Africa. Emergence paths may be combined. For instance, Paths 1 and 3 are combined in the potato late blight epidemic of the 1990s in Western Europe; Paths 1, 2, and 3 are combined in the emergence of stripe rust in Western Europe.

**Fig. 1.**
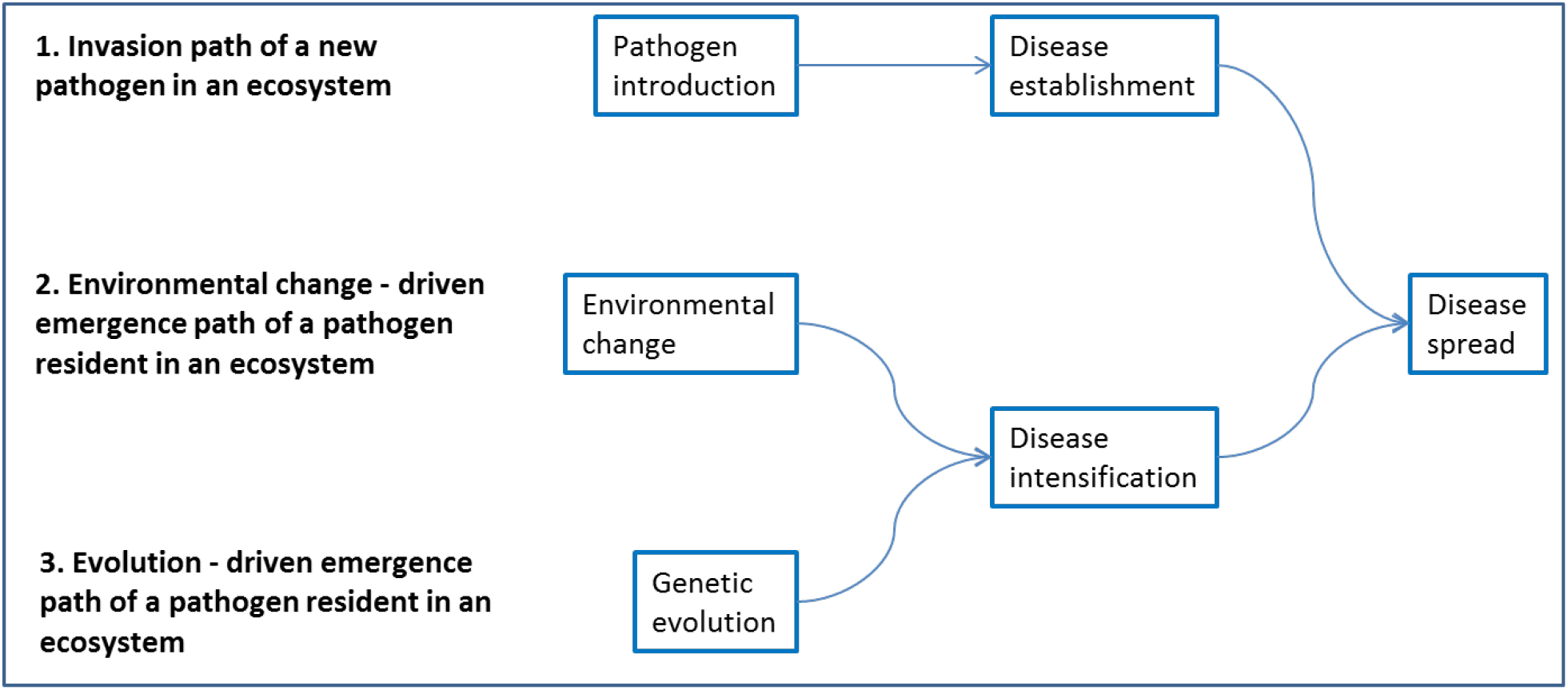
A framework for analysis of emerging epidemics: paths and processes. Three paths for emergence are considered (left, bold characters), involving different processes (in boxes). Paths may be combined, e.g., paths 1 and 3, involving both introduction and evolution, or 2 and 3, involving environmental change and evolution. See text for examples.

Similar to the emergence and re-emergence of infectious diseases in humans (Wilcox and Collwell, 2005), the emergence of plant diseases entails the consideration of biocomplexity, i.e., of complex systems, where the biology of pathogens and hosts, their genetics, the changing environments – both natural and human-made, and the social and economic structures (including plant health management systems) interact. The present analysis does not address the biocomplexity of plant disease emergence as a whole, but rather focuses on a fragment of Figure 1, with emphasis on Paths 1 (emerging pathogens) and 3 (emerging strains). Elements of Path 2 (environmental change) are subsumed in the form of stochastic features of the modelling work.

Here we present a series of hypotheses underpinning the processes at play in disease emergence. These hypotheses involve both demography (epidemiology) and population genetics as follows:

1. from an epidemiological standpoint, emergence is a polyetic process, i.e., it is a process spanning several consecutive crop seasons (Zadoks, 1974; Zadoks and Schein, 1979; Heesterbeek and Zadoks, 1987);
2. this polyetic process is inherently stochastic because it entails random and abrupt changes in the pathogen and host populations (Shaw, 1994). The process is also affected by random fluctuations in the environment (Gilligan and Van den Bosch, 2008);
3. an important determinant of successful emergence is the diversity in the population from which the emerging pathogen originates. We assume the pathogen (or pathogen strain) to be sampled by chance in a large genetic pool. The more diverse this pool, the higher the likelihood of fit to a given biological (hosts) and physical setting (McDonald and Stukenbrock, 2016);
4. pathogen migration (introduction) is often the primary mechanism associated with disease emergence (McDonald and Stukenbrock, 2016);
5. the level of crop losses associated with epidemics constitutes a useful metric for the impact of disease emergence (Savary *et al.*, 2006; 2017; 2019).

A range of models have been developed to analyse the dynamics of epidemics or pathogen populations over multiple crop seasons. Leonard (1977) analysed the dynamics of plant pathogen genotypes over seasons to investigate plant pathogen evolution under the gene-for-gene hypothesis. Since then, several polyetic models have considered cycles of epidemic processes (disease transmission in the presence of the host) followed by survival processes (pathogen decay in the absence of the host). Several models have considered one pathogen genotype in order to address, e.g., thresholds for persistence according to epidemiological parameters (e.g., Gubbins *et al.*, 2000; Madden and Van den Bosch, 2002), whereby persistence corresponds to disease emergence caused by invasion. These models were expanded to consider two pathogen genotypes to analyse the evolutionary dynamics of pathogen populations (Van den Berg *et al.*, 2011; Hamelin *et al.*, 2011). Comparatively fewer stochastic polyetic models have been developed, showing chaotic polyetic patterns (Shaw, 1994), or guiding management strategies (with a spatially explicit stochastic model of sugar beet rhizomania; Gilligan *et al.*, 2007). To our knowledge, no model has yet been developed which simultaneously accounts for polyetic processes, stochasticity, and the occurrence of several pathogen genotypes. Furthermore, none of the polyetic models reported so far explicitly accounts for the impact of disease on yield loss.

The objectives of this work were to: (1) design a modelling framework to better define the conditions determining disease emergence, (2) illustrate the use of the model by considering fitness components that characterize the growth of the pathogen population during the growing season and its survival between growing seasons and analysing their effects on disease emergence, and (3) draw some conclusions on properties associated with disease emergence.

## 2. MATERIALS AND METHODS

### 2.1 Model requirements

Model specifications were established to build a structure incorporating processes related to population genetics, epidemiology, and crop losses in order to analyse the conditions associated with disease emergence and its effects on yield. The model and the outcomes of the analysis apply equally well to: (i) the situation when a strain of a new pathogen is emerging on the background of another, resident pathogen population infecting the same host, and (ii) the situation where a new, immigrant strain of a pathogen emerges on the background of an already established population of the same pathogen. For the sake of simplicity, we will only refer to the second situation below. The following requirements for the model were identified:

- Because disease emergence takes place over several crop seasons (Gilligan and Van den Bosch, 2008), the model generates dynamics of epidemics and pathogen survival over successive cropping cycles, i.e., it encapsulates *polyetic processes* (Zadoks, 1974; Zadoks and Schein, 1979; Heesterbeek and Zadoks, 1987).
- Epidemics of many plant diseases entail secondary infections occurring during a crop season. The model therefore considers *polycyclic epidemics* within crop seasons.
- The model involves *different strains of a pathogen* in order to account for the evolutionary processes involved in disease emergence.
- Disease emergence often originates from the *migration* of a new pathogen (Fig. 1, Path 1), or of a new pathogen strain (Fig. 1, Path 3), into an agrosystem. The model therefore incorporates an immigration process.
- Modelling of the dynamics of *primary inoculum* with varying numbers of propagules, originating from preceding crop seasons and/or from immigration, and decaying over time, is a requirement, because (1) primary inoculum enables the initiation of seasonal epidemics, (2) a migrating pathogen strain enters the system as primary inoculum, and (3) primary inoculum also constitutes the link between two seasonal epidemics, and therefore provides the bridge needed to consider polyetic epidemics.
- C*rop losses* are an essential feature of epidemics in agroecosystems. The model therefore translates multi-seasonal, polyetic epidemics into their impact on crop performance as yield losses.

### 2.2 Model description

The system considered in the model is 1 m^2^ of a crop, under the “mean-field” hypothesis: the system considered is surrounded by systems with the same features and dynamics. This 1 m^2^-system and its surroundings are repeated in successive crop seasons separated by off-seasons. During any crop season, this system and its neighbours are considered in a steady-state relationship. In particular, incoming and outgoing inoculum between these systems cancel each other out, so that the net inflow/outflow balance is null. The model time step is one day, so as to accommodate processes which can have fast dynamics, such as polycyclic processes.

We consider a crop that is grown in regular cropping cycles (Supplementary Figure 1). Each cropping cycle consists of the period when the crop is present (crop season) and the period when the crop is absent (off-season). The crop season starts from crop planting and ends at harvest, and has two phases: the crop establishment phase and the crop growth phase (or, shortly, the growing season). The duration of each of the two phases (*CEP*, crop establishment period, and *CGP*, crop growth period) can vary depending on the crop, crop type (winter or spring crop) and location. Simulations start at crop planting, and are run for 30 cropping cycles.

The model considers one host plant genotype, e.g. a variety of a given crop, which can be infected by two strains of a given pathogen: a local (or resident) strain and an exogenous (or immigrant) strain. The local pathogen strain is present at the beginning of the simulation, while the exogenous pathogen strain is introduced into the system during the course of the simulation. The local population consists of strains that are already well adapted to local conditions. This local population is represented in the model by one local strain which has fixed demographic parameters. The exogenous population is established in a range of conditions (outside from the system), which may differ from the conditions of the considered system. This population is therefore more diverse, and generally less well adapted to the local conditions of the system. It thus consists of strains with a broader range of fitness attributes compared to the local population. This exogenous population is represented in the model by one strain with a fitness that can vary over cropping cycles. This variation reflects the hypothesis that the exogenous strain is less well adapted to the local conditions than the resident strain, and therefore is less well adapted to the environmental variations over cropping cycles. Each cropping cycle therefore involves two strains of the pathogen, the (fixed) local strain, and the (variable, random) exogenous strain.

Each cropping cycle involves several processes, which are represented as rates (Forrester, 1961; Savary and Willocquet, 2014) in Figure 2. These are the processes involved in the development of epidemics, including primary and secondary infections: *RI* (rate of infection); processes involved in the survival and decay of inoculum: *Rdecay* (rate of inoculum decay); and processes involved in yield losses incurred from disease: *RL* (rate of loss). These processes are next described in greater detail. The model variables and parameters are described in Table 1.

**Table 1.** Description of the model variables

| Acronym | Definition | Dimension | Unit | Value |
| --- | --- | --- | --- | --- |
| State variables: |  |  |  |  |
| *i* | Injury caused by disease on a crop stand | [-] | % | 0 to 100 |
| *S* | Number of surviving propagules | [-] | % | 0 to 100 |
| *YL* | Yield loss – yield reduction from a disease-free attainable yield | [-] | % | 0 to 100 |
| Rates: |  |  |  |  |
| *RPI* | Rate of primary infections | [T<sup>-1</sup>] | % Day<sup>-1</sup> |  |
| *RconvIS* | Rate of conversion from injury (i) into surviving propagules (S) | [T<sup>-1</sup>] | % Day<sup>-1</sup> |  |
| *Rdecay* | Rate of decay of surviving propagules | [T<sup>-1</sup>] | % Day<sup>-1</sup> |  |
| *resetYL* | Rate of reset of yield loss at each cropping cycle | [T<sup>-1</sup>] | % Day<sup>-1</sup> |  |
| *RI* | Rate of injury increase | [T<sup>-1</sup>] | % Day<sup>-1</sup> |  |
| *RL* | Rate of yield loss | [T<sup>-1</sup>] | % Day<sup>-1</sup> |  |
| *RM* | Rate of immigration of the pathogen | [T<sup>-1</sup>] | % Day<sup>-1</sup> |  |
| *starter* | Rate of infection to initiate epidemics | [T<sup>-1</sup>] | % Day<sup>-1</sup> |  |
| Parameters: |  |  |  |  |
| *CEP* | Duration of the crop establishment period | [T] | Day | 120 |
| *CGP* | Duration of the crop growth period | [T] | Day | 120 |
| *convSP* | Conversion of surviving propagules into a rate of primary infections | [T<sup>-1</sup>] | Day<sup>-1</sup> | 0.01 |
| *imax* | Carrying capacity of injury – maximum level of injury | [-] | % | 100 |
| *RRD<sub>j</sub>* | Relative rate of decay of surviving propagules for strain j | [T<sup>-1</sup>] | Day<sup>-1</sup> | j = 1: 0.01<br>j = 2: Varying |
| *RRg<sub>j</sub>* | Relative rate of epidemic (injury) increase for strain j | [T<sup>-1</sup>] | Day<sup>-1</sup> | j = 1: 0.07<br>j = 2: Varying |
| *RRL* | Relative rate of yield loss | [T<sup>-1</sup>] | Day<sup>-1</sup> | 0.05 |
| *Ya* | Attainable yield – uninjured yield level | [-] | % | 100 |

**Fig. 2.**
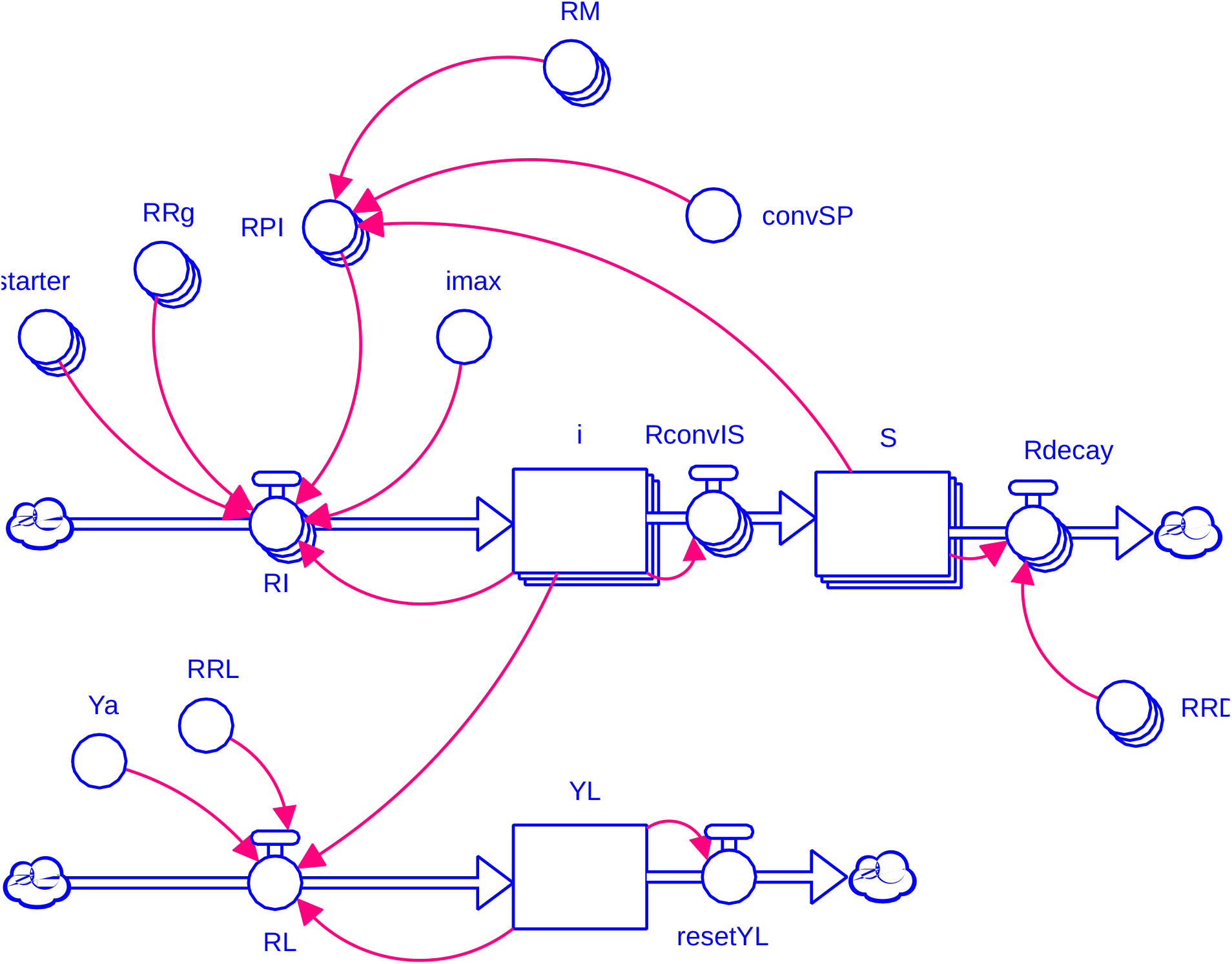
Simplified flowchart of the process-based model used to analyse emerging diseases of crop plants. Variable acronyms are described in Table 1. The flowchart uses symbols introduced by J Forrester (Forrester, 1961): rectangles represent state variables; valves represent rates of change of state variables; circles represent parameters or computed variables. Stacked symbols (e.g., state variables) represent vectors of two pathogen strains.

In each cropping cycle, the epidemic starts with primary infections (*RPI*), which take place at the end of the crop establishment phase, as the crop growth phase starts. Primary infections have two origins. First, primary infections can originate from inoculum produced from epidemics which took place in previous crop seasons (polyetism), and second, primary infections can result from incoming inoculum (immigration from an exogenous population). In the beginning of the crop growth phase, the rate of primary infections for each strain, *j* = 1 (local), or 2 (immigrant), is therefore written as:

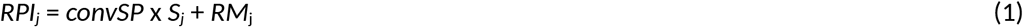

Where *convSP* is the conversion of surviving inoculum into a rate of primary infections; *S*_j_ is the number of surviving propagules for each strain; and *RM*_j_ is the rate of infections originating from immigrant strains, referred to as the rate of immigration. The rate of primary infections, *RPI*_j_, has the value given by Eq. (1) only on the first time step of each growing season and is set to zero at all other times.

An epidemic takes place as the injury level, *i*, increases according to a logistic curve (exponential increase of secondary infections, limited by the carrying capacity of the host crop) with a relative rate of growth, *RRg.* As the seasonal epidemic unfolds, interaction between strains takes place, in the form of competition towards host (crop) sites. This interaction between strains accounts for the maximum possible level of injury (carrying capacity) at a given time, considering all plant sites occupied by the different strains at this time. The rate of infection of each strain *j*, comprising primary and secondary infections, is therefore written as:

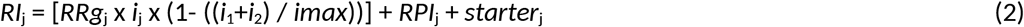

where *RRg*_j_ is the relative rate of injury increase for strain *j*; *i*_j_ is the injury level of strain *j*; *i*_1_ is the injury level caused by the local strain; *i*_2_ is the injury level caused by the immigrant strain; *imax* is the carrying capacity of injury, i.e., the maximum level of injury; *RPI*_j_ is the rate of primary infections associated with strain *j*; and *starter*_j_ is the number of primary infections at the beginning of the multiple-cropping cycle simulation (this parameter is non-zero only during the first time step of the cropping cycle 1).

At the end of a cropping cycle, the terminal injury level (*i*_j_) is converted into surviving inoculum, *S*_j_, for each of the two strains. The number of surviving propagules decreases over time according to a negative exponential dynamics, at a speed proportional to a relative rate of decay (*RRD*_j_):

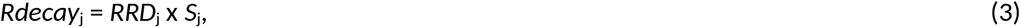

where *Rdecay*_j_ is the rate of decay of surviving propagules of strain *j*; *RRD*_j_ is the relative rate of decay of surviving propagules of strain *j*, and *S*_j_ is the number of surviving propagules of strain *j.*

Injuries impair the physiological processes involved in crop growth and yield build-up, ultimately leading to yield losses. The several possible damage mechanisms from injuries are represented in a very simplified manner by a single rate of yield loss, *RL*, which increases proportionally to the running level of combined injuries caused by both strains, *i*_1_ + *i*_2_:

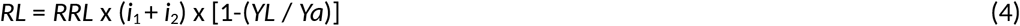

where *RRL* is the relative rate of yield loss; *i*_1_ and *i*_2_ are the injury levels from the local and immigrant strains, respectively; *YL* is the yield loss, i.e. the yield reduction from a disease-free attainable yield; and *Ya* is the attainable yield, i.e., the yield level in the absence of disease. At the end of each crop growth phase, yield loss is reset to zero, so that the new cropping cycle starts without losses.

### 2.3 Model parameters and initial values

Initial values are zero for all state variables (*i*, *S*, and *YL*). Parameters dimensions and values are listed in Table 1. The durations of the crop establishment period (*CEP*) and of the crop growth period (*CGP*) are both set to 120 days, representing, for example, approximate durations for a winter wheat crop grown in a temperate region of the world. An epidemic of the local strain is initiated at the end of *CEP* in the first cropping cycle with an initial value (*starter*; Table 1) for the local strain of 0.01 day^−1^. The conversion of surviving propagules into a rate of primary infections (*convSP*) is set to 0.01, meaning that for example 100 surviving entities are translated into 1 primary infection during the first time step of CGP. The carrying capacity for injury level, *imax*, is set to 100 in order to generate injury levels expressed as percent. In the same way, *Ya*, the attainable yield, is set to 100 in order to generate yield losses expressed as percent.

*RRL* is set to 0.05, meaning that combined disease injury (*i*_1_ + *i*_2_), when at low levels, entails an increase in yield loss at each time step which corresponds to 5% of the level of disease injury. *RRg*_1_ and *RRD*_1_ values are set to 0.07 and 0.01, respectively.

### 2.4 Model analyses: conditions of emergence of an immigrant strain

#### 2.4.1 Framework of analyses

We consider a pathosystem with two pathogen strains: a local (resident), and an immigrant (exogenous) strain. The fitness of each of the two strains is represented by two essential components: the ability to reproduce during the growing season [represented by a relative, or intrinsic, rate of growth, *RRg*_*j*_ in Equation (2)] and the rate of population decay [represented by a relative, or intrinsic, rate of decay, *RRD*_*j*_ in Equation (3), Table 1], the latter characterizing the ability of a pathogen strain to survive in the absence of host plants. As a convention, the subscript *j*=1 refers to a local strain and *j*=2 refers to an immigrant strain. Fitter strains reproduce faster on the host during the growing season and decay more slowly over time.

We addressed the question of emergence of immigrant strains as follows. A given agroecosystem harbours a resident diversity of strains; however, all these strains are assumed to be equally adapted to the considered agroecosystem – i.e. they have similar fitness. As a simplification, the entire population of resident strains in an agroecosystem is therefore represented by one strain, exhibiting two central values for *RRg*_1_ and *RRD*_1_. Because these local populations are assumed to be established and in a dynamic equilibrium, we further assume no variation over time for parameters *RRg*_1_ and *RRD*_1_. In the absence of immigration, successive epidemics occur in the considered agroecosystem. These epidemics consist of overlapping disease cycles (polycyclic epidemics), and each epidemic results from the carry-over of inoculum from a previous epidemic that took place in the previous crop seasons. The resulting pattern of disease over successive crop seasons (polyetic process) in an agroecosystem thus results from the concatenation of successive (polycyclic) epidemics.

In order to investigate conditions for emergence, we consider an immigrant strain, which originates from a very large pool of possible strains. In a first (deterministic) regime, the fitness parameters of the immigrant strain, *RRg*_*2*_ and *RRD*_*2*_, are assumed to be constant throughout the successive simulated cropping seasons. In a second (stochastic) regime, the fitness parameters of the immigrant strain are drawn at random from a normal distribution with central values *RRg*_*2*_ and *RRD*_*2*_, and with variation about these values. This drawing is made at the beginning of each cropping cycle, and the values drawn are kept constant within each cropping cycle. This stochastic regime reflects the hypothesis of a strain which is not well adapted to the local environment, with a fitness that varies as environmental conditions vary over cropping cycles.

The execution of the model over a succession of 30 cropping cycles is referred to as a simulation. We investigated a scenario in which the immigrant strain is introduced once, at cropping cycle 10, at the beginning of the growing season. This way, the immigrant strain is introduced into a stabilized system where the local strain is already established. We used the simulation model (Section 2.2, Figure 2) to study two dynamic regimes: (i) a deterministic regime, in which *RRg*_*2*_ and *RRD*_*2*_ had fixed values during a given simulation (section below), and (ii) a stochastic regime, in which the values of either *RRg*_2_ or *RRD*_2_ or both were drawn from a normal (Gaussian) distribution at the beginning of each cropping cycle and kept at these values during each cropping cycle (section 2.4.3 below). We also derived approximate analytical expressions for the thresholds of emergence of the immigrant strain by representing the simulation model as a discrete time map and investigating its linear stability (Section 2.4.4 below and Appendix A).

The outcomes of the analyses were synthesised according to three features characterising disease emergence of an immigrant strain and its consequences: the probability of emergence, the time to emergence, and the yield loss associated with the emergence. We consider that the immigrant strain has emerged if it exceeds the resident strain in terms of its AUDPC (area under disease progress curve, i.e., the accumulated injury incurred within a growing season) during at least three cropping cycles after its introduction. The probability of emergence, *P*_emerg_, was estimated as the proportion of simulations that resulted in emergence. In each individual simulation that resulted in emergence, the time to emergence, *T*_emerg_, was defined as the number of cropping cycles between the introduction of the immigrant strain and the first cropping cycle when the AUDPC of the immigrant strain exceeded that of the resident strain. To quantify yield loss in each simulation, we calculated the average yield losses caused by both the resident and the immigrant pathogen strains over the 30 cropping cycles.

The model was developed using the Stella software (STELLA Architect version 1.1.2) and subsequently translated to the Python programming language (version 3.4.3), where the bulk of the analysis was conducted. The system of Equations (1)-(4) was solved and analysed using Python packages numpy (version 1.13.3) and scipy (version 1.0.0), and the figures were produced using the Python package matplotlib (version 2.1.1). Parts of the analytical investigation were performed with Wolfram Mathematica (version 10.3 for Linux).

#### 2.4.2. Deterministic approach

We performed three sets of simulations in order to analyse the individual effects of *RRg*_*2*_, *RRD*_*2*_, and the combined effects of *RRg*_*2*_ and *RRD*_*2*_ on disease emergence.

A first analysis was conducted to address conditions of emergence associated to *RRg*_*2*_. In this first analysis, 100 simulations were run with *RRg*_*2*_ increasing from 0.06 to 0.12 day^−1^ with a constant increment of *RRg*_*2*_ between simulations, while *RRD*_*2*_ was fixed (0.02 day^−1^). The *RRD*_*2*_ value chosen corresponds to the hypothesis of an immigrant strain with a lower survival capacity than the resident strain (*RRD*_*1*_ = 0.01 day^−1^). In the second analysis, we assessed conditions of emergence according to *RRD*_*2*_. Here, 100 simulations were run with *RRD*_*2*_ increasing from 0 to 0.05 day^−1^ with a constant increment between simulations, while *RRg*_*2*_ was fixed (0.1 day^−1^). This *RRg*_*2*_ value corresponds to the hypothesis of an immigrant strain with a higher aggressiveness than the resident strain (*RRg*_*1*_ = 0.07 day^−1^). In a third analysis, both *RRg*_*2*_ and *RRD*_*2*_ were considered with respect to emergence. *RRg*_*2*_ and *RRD*_*2*_ were varied in the same ranges as in the first and second analyses over a total of 10^4^ simulations (100 × 100 runs).

#### 2.4.3. Stochastic approach

As in the deterministic approach, the individual effects of *RRg*_*2*_, *RRD*_*2*_, and combined effects of *RRg*_*2*_ and *RRD*_*2*_ were subsequently analysed.

To address conditions of emergence associated to *RRg*_*2*_, 100 sets of simulations were executed with fixed *RRg*_2_ values ranging from 0.06 to 0.12 day^−1^, with a constant increment. For each *RRg*_*2*_ value considered (i.e., for each set of simulations), 5000 stochastic runs were executed, within which the values of *RRD*_2_ were drawn at the beginning of each cropping cycle as random numbers from the normal (Gaussian) distribution with the mean 0.02 day^−1^ and the standard deviation 0.007 day^−1^. These *RRD*_2_ values then remained constant during the whole cropping cycle until the beginning of the next growing season, when a new random value was chosen.

The second analysis was conducted in the same way as the first analysis, but focused on *RRD*_*2*_: 100 sets of simulations were executed with fixed *RRD*_2_ values ranging from 0 to 0.05 day^−1^, with a constant increment. For each *RRD*_2_ value considered, 5000 stochastic runs were executed, within which the values of *RRg*_2_ were drawn at the beginning of each cropping cycle as random numbers from the normal distribution with mean 0.1 day^−1^ and standard deviation 0.035 day^−1^. These *RRg*_2_ values then remained constant during the whole cropping cycle.

In a third analysis, the values of both *RRg*_*2*_ and *RRD*_*2*_ were drawn from the normal distribution at the beginning of each cropping cycle with means ranging from 0.05 to 0.12 day^−1^ for *RRG*_*2*_, and ranging from 0 to 0.04 for *RRD*_*2*_, and with standard deviations constituting a constant proportion, 0.35, of the corresponding mean values. As in the previous analyses, *RRg*_1_ and *RRD*_1_ values remained constant within each cropping cycle. We ran 200 stochastic realizations for each point of the 100×100 grid of *RRg*_2_ × *RRD*_2_ values considered.

#### 2.4.4. Analytical approach

The overall fitness of the pathogen strain *j* is given by its polyetic (or multi-season) basic reproductive number (see Appendix A for the derivation):

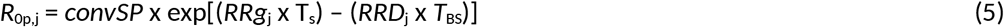

where *T*_S_ is the crop growth duration (*T*_S_ = *CGP*) and *T*_BS_ is the delay between two successive growing seasons. The index “p” in *R*_0p_ refers to “polyetic”, in order to distinguish *R*_0p_ from *R*_0_, which usually refers to the “within season” basic reproductive number in the epidemiological literature (e.g., Zadoks and Schein, 1979; Anderson and May, 1986; Campbell and Madden, 1990). Biologically, *R*_0p,j_ represents the number of units of crop injury appearing at the beginning of a given growing season following the introduction of a unit host injury in the beginning of the previous growing season. *R*_0p,j_ incorporates both the ability of a strain *j* to multiply during crop growth and to survive between growing seasons. Hence, in the exponent of Eq. (5), the two components of pathogen fitness, *RRg*_*j*_ and *RRD*_*j*_, are weighted by *T*_S_ and *T*_BS_, respectively.

As in the previous analysis, we consider the situation when the local pathogen strain is viable when present alone: its reproduction during the growing season exceeds its losses between growing seasons, i.e., *R*_0p,1_>1. In this case, the immigrant strain will emerge if its polyetic basic reproductive number exceeds the polyetic basic reproductive number of the local strain, i.e., *R*_0p,2_>*R*_0p,1_. The emergence threshold corresponds to *R*_0p,2_ = *R*_0p,1_. We solve this equation with respect to *RRg*_2_ and obtain the threshold value of *RRg*_2_ above which emergence takes place:

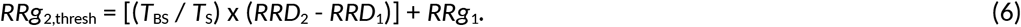

Similarly, the emergence threshold can be expressed in terms of *RRD*_*2*_:

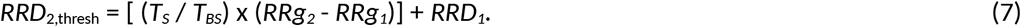

Here, the immigrant strain emerges when its relative rate of decay is below the threshold, i.e., *RRD*_2_<*RRD*_2,thresh_. Eqs. (6) and (7) were derived under the assumption that the saturation effects of the logistic growth are negligible. This is justified at sufficiently low relative rates of growth for each of the strains *RRg*_*j*_, and at short enough *CGP*, so that the host tissue does not become a limiting factor for any of the two pathogen strains. Note that the simulation model in Sec. 2.2 does not make this approximation. See Appendix A for more mathematical details.

## 3. RESULTS

### 3.1 An example of dynamics of crop injuries and losses simulated with the stochastic approach

Figure 3 displays examples of simulated dynamics using fitness parameters for the immigrant strain drawn from a normal distribution at the beginning of each cropping cycle, the parameter values remaining fixed within a given cropping cycle. Means of *RRg*_2_ and *RRD*_2_ are equal to the values used for the local strain (0.07 for *RRg* and 0.01 for *RRD*) and their standard deviations are 0.03 and 0.003, respectively. The three top panels display injury dynamics leading to (1) non-emergence of the immigrant strain (Fig. 3a), (2) co-occurrence of both strains where the predominant strain varies over cropping cycles (Fig. 3b), and (3) rapid emergence of the immigrant strain (Fig. 3c). Because parameters are fixed for the local strain, simulation leading to non-emergence (Fig. 3a) shows an equilibrium state with a maximum level of injury which reaches a constant value starting from the 6^th^ cropping cycle. When both strains co-exist (Fig. 3b), the stochasticity of *RRg*_2_ and *RRD*_2_ produces a large variation over cropping cycles in both injury level and the respective frequency of each strain. In the case of rapid emergence of the immigrant strain (Fig. 3c), the immigrant strain very quickly overcomes the local strain, but displays a large variation in disease intensity over cropping cycles, because of stochasticity in *RRg*_2_ and *RRD*_2_.

**Fig. 3.**
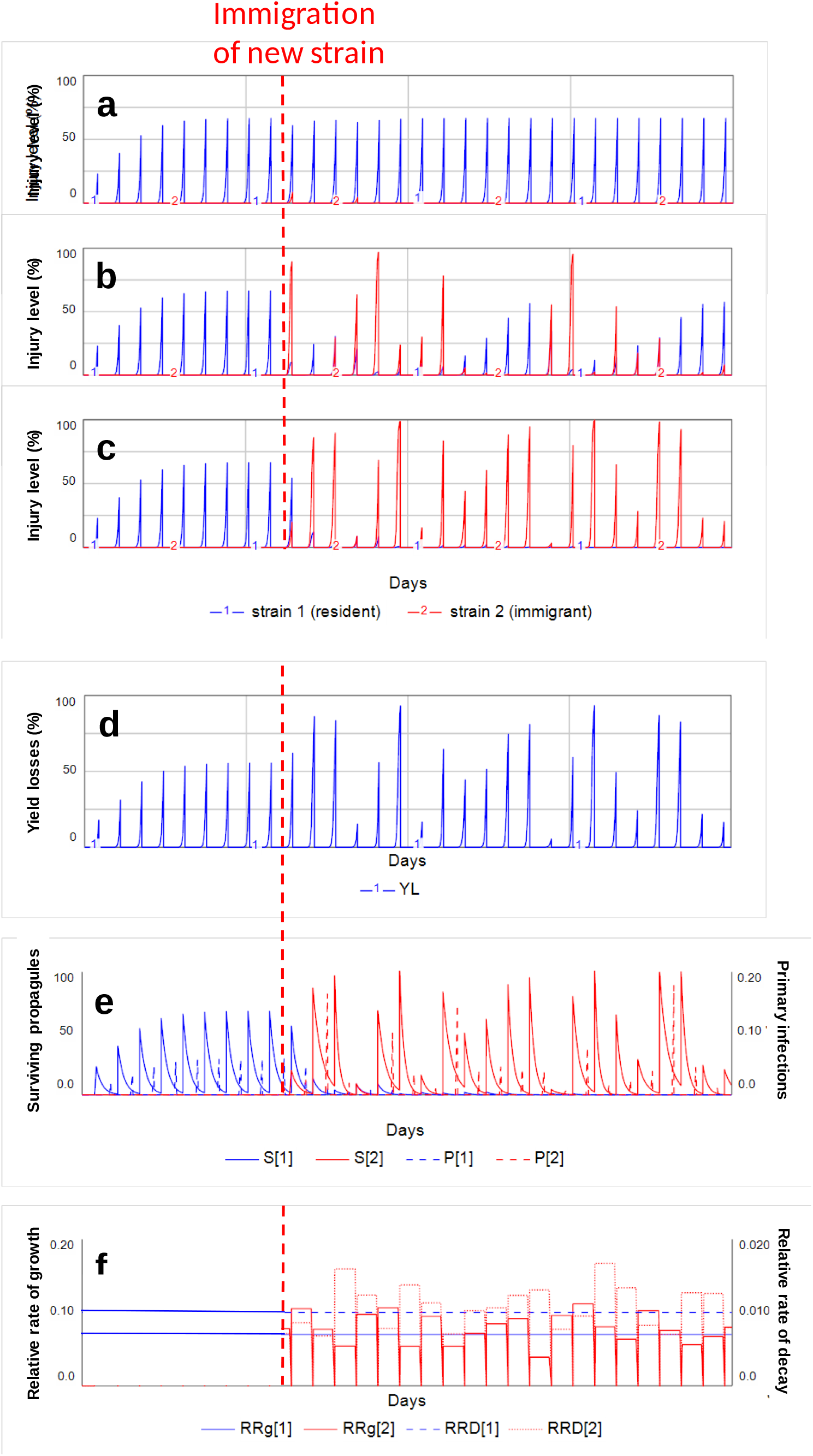
Examples of simulated dynamics of injury levels and related variables over 30 cropping cycles. (a) to (c): simulated injury levels from different runs where *RRg*_2_~N(0.07, 0.03) and *RRD*_2_~N(0.01, 0003). (a): a simulation where the immigrant strain did not emerge; (b): a simulation where the immigrant and resident strains compete over cropping cycles; (c): a simulation where the immigrant strain replaces the resident strain; (d) to (f): simulated dynamics of other variables, associated with the dynamic of injury levels shown in (c). Except for *RRg*_2_ and *RRD*_2_, all parameters are set according to Table 1. Immigration of strain 2 takes place at the end of the crop establishment period of cropping cycle 10, while strain 1 is established at the beginning of the simulation.

Simulated yield loss, primary inoculum, and relative rates of growth and decay corresponding to the example of a rapidly emerging strain (Fig. 3c) are displayed in Figure 3d-f. Yield losses vary over time (Fig. 3d), with a pattern similar to that observed for injury dynamics (Fig. 3c). At the end of each crop season, the terminal disease injury from each strain is proportionally converted to surviving propagules. The number of surviving propagules then decays exponentially over time and constitutes the primary inoculum for the subsequent growing season (Fig. 3e). This primary inoculum translates into primary infections at the beginning of each crop growth phase (Fig. 3e). Relative rates of growth and of decay of the immigrant strain vary over cropping cycles, while remaining constant within each cropping cycle (Fig. 3e). These stochastic values of *RRg*_2_ and *RRD*_2_ are driving the dynamics of injury (Fig. 3c) and of primary inoculum (Fig. 3e) over cropping cycles.

### 3.2 Individual effects of the relative rates of epidemic growth and inoculum decay on disease emergence

When we consider *RRg*_2_ variation in the deterministic regime, *P*_emerg_ rises suddenly from zero to one as the *RRg*_2_ value is increased (Fig. 4a, blue curve). The reason is that the immigrant strain can only emerge if it is able to grow fast enough during the growing season. More fit immigrants, when they emerge (i.e., when *P*_*emerg*_ = 1 in Fig. 4a), emerge more rapidly as *RRg*_*2*_ increases: *T*_emerg_ decreases monotonically as we increase the immigrant strain’s fitness by increasing *RRg*_2_ (Fig. 4c, blue curve). As we increase *RRg*_2_, the amount of disease caused by the immigrant strain increases and so does the average yield losses incurred by both the resident and the immigrant pathogen strains (Fig. 4e, blue curve). Below the emergence threshold (Fig. 4a, *RRg*_2_ values for which *P*_emerg_ = 0), the immigrant strain is absent, therefore the yield losses are only incurred by the resident strain, and are not affected by *RRg*_2_. The analytical approach yields a threshold for emergence of *RRg*_2_ at 0.09 day^−1^ (Fig. 4a, grey line), that is, slightly smaller than the threshold derived from the deterministic approach (Fig. 4a, blue line).

**Fig. 4.**
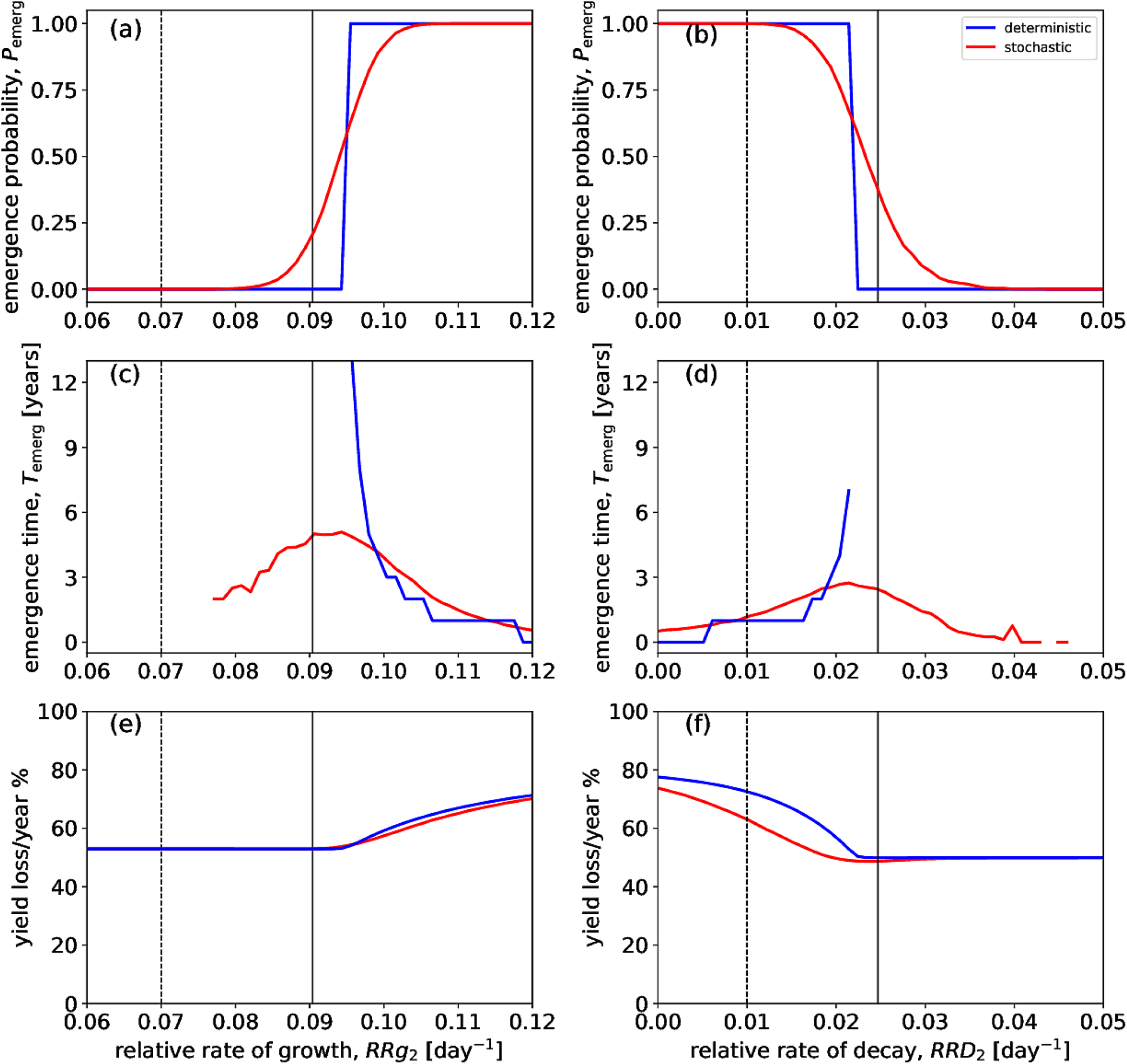
Effects of pathogen reproduction and survival parameters on disease emergence and yield loss. Emergence probabilities (a), (b), emergence times (c), (d) and yield losses (e), (f) are plotted versus the relative rate of growth, *RRg*_2_ (a), (c), (e), and the relative rate of decay, *RRD*_2_ (b), (d), (f), of the immigrant strain. Blue curves correspond to the deterministic regime, in which both *RRg*_2_ and *RRD*_2_ have fixed, deterministic values, whereas red curves correspond to the stochastic regime. Red curves in panels (a), (c), (e) were computed with the values of *RRD*_2_ drawn as random numbers from the normal (Gaussian) distribution at the beginning of each cropping cycle with the mean 0.02 day^−1^ and the standard deviation 0.007 day^−1^, while *RRg*_2_ assumed fixed, deterministic values. Similarly, red curves in panels (b), (d), (f) were computed with the values of *RRg*_2_ drawn as random numbers from the normal (Gaussian) distribution at the beginning of each cropping cycle with the mean 0.1 day^−1^ and the standard deviation 0.035 day^−1^, while *RRD*_2_ assumed fixed, deterministic values. Dashed vertical lines indicate the value of the relative rate of growth, *RRg*_1_=0.07, of the resident strain in (a), (c), (e), and the value of the relative rate of decay, *RRD*_1_=0.01, of the resident strain in (b), (d), (f). Solid vertical lines show the approximate analytical emergence thresholds, according to Eq. (6) in (a), (c), (e) and Eq. (7) in (b), (d), (f). Values of other parameters are given in Table 1.

When we include stochasticity, the transition between parameter areas of “no emergence” and “emergence” is now gradual: *P*_emerg_ increases continuously as *RRg*_2_ values of the immigrant strain are increased (red curve in Fig. 4a). Time to emergence also exhibits a different pattern in the stochastic regime. Emergence starts at *RRg*_2_ values much smaller than in the deterministic regime (Fig. 4c, red curve) with relatively small values of *T*_emerg_ (about 2 cropping cycles), then *T*_emerg_ increases gradually, plateaus at about four cropping cycles, and then declines with a curve close to, but above, that generated from the deterministic approach. Yield losses show a similar pattern in the deterministic and stochastic regimes (compare red and blue curves in Fig. 4e), although yield losses are somewhat lower in the stochastic regime than in the deterministic regime.

The effect of *RRD*_2_ on emergence characteristics (Fig. 4b, d, f) mirrors the effect of *RRg*_2_ (Fig. 4a, c, e), because as fitness of the immigrant strain increases with *RRg*_2_, it decreases with *RRD*_2_. Under the deterministic regime, *P*_emerg_ drops abruptly from one to zero as *RRD*_2_ is increased (Fig. 4b, blue curve): the immigrant strain cannot emerge if its population decays too fast between growing seasons. In the same way, less fit immigrants emerge more slowly, when they do emerge: *T*_emerg_ increases monotonically as we reduce the immigrant strain’s fitness as *RRD*_2_ increases (Fig. 4d, blue line). Average yield losses decrease as *RRD*_2_ increases (Fig. 4f), and remain stable when *RRD*_2_ values are above the threshold for emergence. The threshold for emergence generated from the analytical approach, *RRD*_2_ = 0.025, is slightly lower than the threshold generated from the deterministic approach (Fig. 4b).

When we include stochasticity, *P*_emerg_ diminishes continuously as *RRD*_2_ is increased; time to emergence, *T*_emerg_, increases initially (with values slightly larger than those obtained from the deterministic approach), reaches a maximum around the emergence threshold and gradually declines to small values. Yield losses show a qualitatively similar pattern in the deterministic and stochastic regimes. As when investigating the effect of *RRg*_*2*_, yield losses are lower in the stochastic regime than in the deterministic regime (Fig. 4f).

### 3.3 Combined effects of the relative rates of epidemic growth and inoculum decay on disease emergence

When using the deterministic approach, emergence and no emergence domains are clearly separated by a straight line (Fig. 5a). This line reflects the abrupt transition from emergence to no emergence, as illustrated in Figs. 4a and 4d. The domain of emergence corresponds to pairs of values of *RRg*_*2*_ and *RRD*_*2*_ below which emergence takes place: for a given value of *RRg*_*2*_, emergence will occur within a range of values of *RRD*_*2*_ below a given threshold. The dashed lines in Fig. 5a display the outcomes for fitness values used which correspond to that of the resident strain. At *RRg*_*2*_ = *RRg*_*1*_ = 0.07, emergence occurs for *RRD*_*2*_ values slightly smaller than *RRD*_*1*_. Similarly, at *RRD*_*2*_ = *RRD*_*1*_ = 0.01, emergence occurs for *RRg*_*2*_ values that are slightly larger than *RRg*_*1*_. The solid grey line represents the analytical expression for the emergence threshold in terms of *RRD*_*2*_, according to Equation (7). That is, a line with slope (*T*_*S*_ / *T*_*BS*_) which equals 0.5 in our case, and an ordinate at origin of −0.025.

**Fig. 5.**
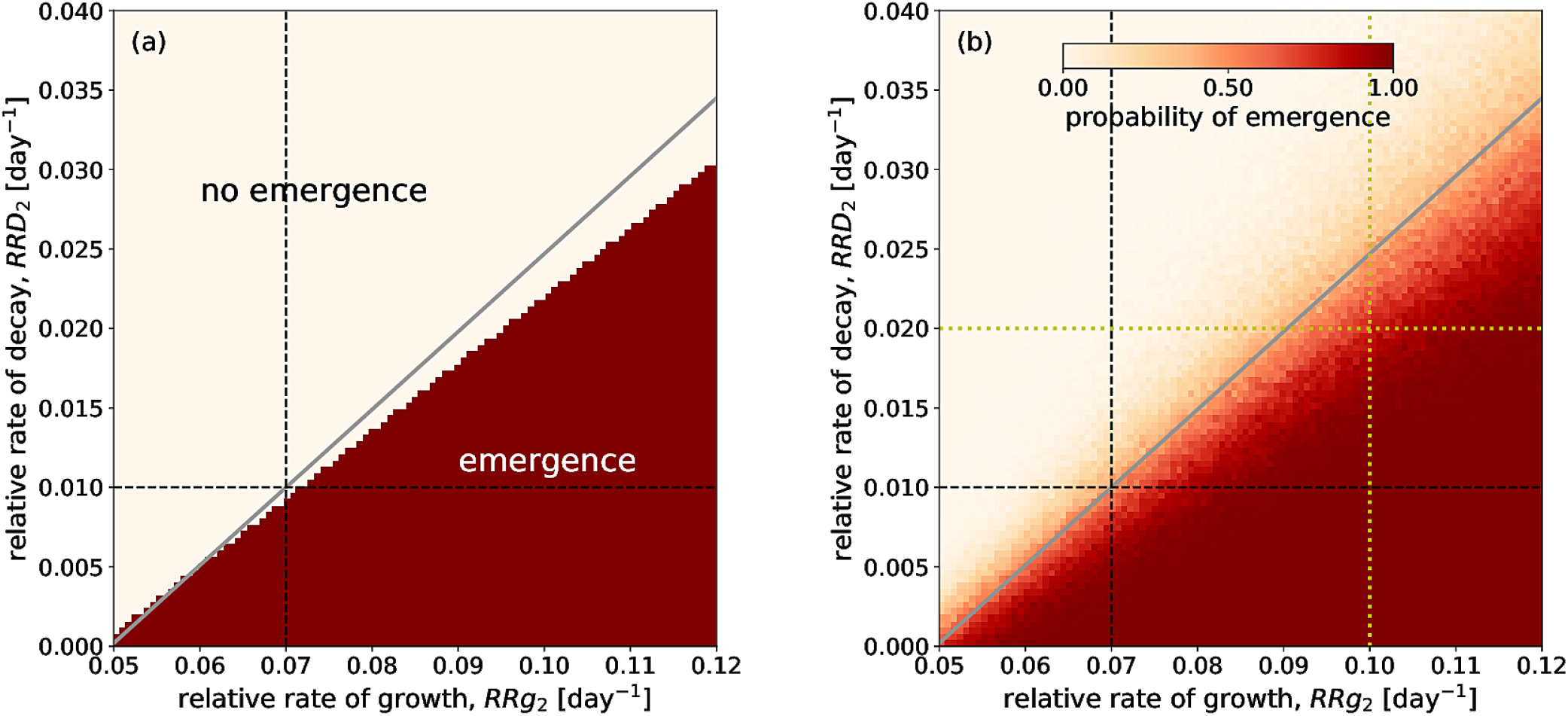
Combined effects of pathogen reproduction and survival parameters on probability of emergence. (a) deterministic regime; (b) stochastic regime, whereby the values of *RRg*_2_ and *RRD*_2_ were drawn from the normal (Gaussian) distribution at the beginning of each cropping cycle with the mean corresponding to the values on x- and y-axes and the standard deviation constituting a constant proportion, 0.35, of the corresponding mean values. Values of other parameters are given in Table 1. In each panel, the grey diagonal line shows the analytical threshold according to Eq. (7). Dashed black lines indicate the fitness values of the resident strain: vertical *RRg*_1_=0.07 day^−1^; horizontal *RRD*_1_=0.01 day^−1^. Dashed yellow lines mark the parameter regions explored in Fig. 4: *RRD*_2_=0.02 day^−1^ in Fig. 4a, c, e; *RRg*_2_=0.1 day^−1^ in Fig. 4b, d, f.

When stochasticity is included in the model, the transition between parameter domains of emergence and no emergence becomes gradual (Fig. 5b). This gradual transition is a generalisation of the gradual change in emergence probability according to *RRg*_2_ and *RRD*_*2*_ illustrated in Figs. 4a and 4d, respectively. As in Fig. 5a, the dashed black line represents the fitness parameters of the resident stain. In the same way, the parameter region explored in Fig. 4a and 4b is materialised with the yellow lines which refer to *RRg*_*2*_ = 0.1 and *RRD*_*2*_ = 0.02. The most important effect of stochasticity is that emergence occurs at ranges of parameters that are below the emergence threshold, where no emergence was possible according to the deterministic approach (within the white area in Fig. 5a). Even if the immigrant strain is on average less fit than the local strain (i.e., in terms of its average fitness components *RRg*_2_ < *RRg*_1_, *RRD*_2_ > *RRD*_1_), there is still a non-zero probability for its emergence.

This scenario corresponds to the region in Fig. 5b above the black horizontal line and to the left from the black vertical line.

## 4. DISCUSSION

### 4.1 Key findings

The modelling framework which was designed in this work enables analysing the conditions underlying disease emergence. According to our pre-set specifications, the model includes polyetism, stochasticity, and yield loss. The analyses conducted here allow identifying important features associated with disease emergence.

A major finding is that stochasticity can be an important boundary condition for disease emergence. Emergence reflects important changes in the status of a system (here crop health) that can be caused by rare events (e.g., Paini *et al.*, 2016), associated with small sizes of immigrant or mutant subpopulations that initiate the process, and by polyetic processes that lead to significant reductions in population size between growing seasons (Shaw, 1994). Stochasticity associated with genetic factors such as bottlenecks and genetic drift is known to play an important role in the evolution of a host-pathogen interaction (McDonald, 2004). Stochasticity can also be introduced by environmental factors such as climatic conditions, which can differentially affect the fitness of strains in pathogen populations (e.g., Gilligan and Van den Bosch, 2008). In the present study, we focused on the latter, environmentally-induced, stochasticity.

Previous modelling analyses considered polyetic processes (e.g., Gubbins *et al.*, 2000; Madden and Van den Bosch, 2002) under a deterministic framework, leading to identification of thresholds for persistence. Results generated from the stochastic approach in this work produce a different outcome, in showing that (1) even when an immigrant strain is drawn from a population which is, on average, less fit than the local strain, the immigrant strain may nevertheless emerge due to stochasticity; and conversely (2) even when an immigrant strain is drawn from a population which is, on average, more fit than the local strain, the immigrant strain will not necessarily emerge, and may face extinction. An important implication of this finding is that emergence may require a series of independent immigration events involving new pathogen strains before a successful invasion takes place. In some cases, a pathogen that appears to have suddenly emerged over the course of only 1-2 cropping cycles may have been present at a low level for decades before the proper conditions (e.g. conducive weather conditions) occurred to enable an explosion in population size and an observed “emergence”. This has important implications to guide future research, both in terms of modelling and experimentation, and potentially to inform policy on emerging diseases.

Another important finding from our analyses is that survival between growing seasons is as important for emergence as the pathogen reproduction during the growing season. Pathogens with limited saprophytic abilities and lacking durable survival structures such as chlamydospores, sclerotia or oospores are expected to undergo large bottlenecks between host growing seasons that will purge genetic diversity and increase the probability that less well-adapted immigrants occurring at lower frequencies will go extinct between growing seasons. Conversely, pathogens that compete well as saprophytes and/or produce long-lived survival structures will maintain high effective population sizes that sustain high levels of genetic diversity across growing seasons, enabling persistence of immigrants and novel mutants for long periods of time, even if they are less well adapted, and increasing the probability that these immigrants, mutants or recombinants can make a successful invasion. Although the importance of the survival phase has been recognized in earlier work (e.g., Heesterbeek and Zadoks, 1987; Gubbins *et al.*, 2000; Madden and Van den Bosch, 2002, Hamelin *et al.*, 2011), survival has often been overlooked by plant pathologists. Conversely, *RRg* can be seen as the apparent infection rate of Van der Plank (*r*_*L*_; Campbell and Madden, 1990), for which ranges have been measured from disease progress curves in many instances. The *RRg* ranges explored in our analyses (0.05 to 0.12) fit well within ranges measured for epidemics of annual crop diseases (Kranz, 2003).

### 4.2 Comparing outcomes from the deterministic, analytical, and stochastic approaches

There is a good agreement between the analytical emergence thresholds (Eq. (6) and (7)) and the numerical thresholds in the deterministic regime, although the threshold for emergence with respect to *RRg* is slightly lower when derived from the analytical approach as compared to the deterministic approach (Figs. 4, 5a), while the opposite pattern is obtained for *RRD* (Figs. 4b, 5a). This difference can be explained by the limited duration (30 cropping cycles) of the numerical simulations. In some cases, the immigrant strain would be able to emerge, but this would require more than 30 cropping cycles. On the contrary, the analytical threshold does not restrict the number of cropping cycles and therefore generates thresholds for emergence that can occur over an infinite time span. This explanation was confirmed by performing additional simulations conducted using the same design that generated Figure 4, but including many additional cropping cycles (200). In that case, the agreement between the two thresholds (from deterministic simulations and from analytical expressions) was perfect. In future analyses using this framework, the threshold values to consider (from deterministic or analytical approaches) will depend on the modelling objectives and the applications under consideration.

When investigating the probability of emergence, fitness thresholds are derived from the deterministic approach, while such thresholds do not materialize in the stochastic approach because the emergence probability can take values between 0 and 1. The stochastic approach allows a strain with a fitness (*RRg* or *RRD*) mean value below the deterministic emergence threshold (i.e., lower values for *RRg* and larger values for *RRD*) to emerge, with a probability which progressively declines as the mean fitness value moves away from the threshold.

The time to emergence progressively declines as the fitness values increase in the deterministic approach because it requires progressively less time for the immigrant strain to outcompete the resident strain. Under the stochastic regime, a different pattern is exhibited, with the time for emergence increasing, reaching a maximum, and eventually declining as the fitness values increase. This pattern can be interpreted as follows: at low average fitness, the only way to achieve emergence in the stochastic regime is when high fitness values from the tail of the distribution are drawn over several consecutive growing seasons, representing particularly “lucky” realizations. There is a small proportion of such realizations (reflected by the small probability of emergence), as they correspond to quite rare events, but when they do happen, emergence occurs relatively fast. In contrast, at higher average fitness, there can be many other paths to emergence including those realizations in which high fitness values appear sporadically, not necessarily in several consecutive seasons, leading to slower emergence on average. Thus, the two competing effects, (i) longer emergence due to reduced mean fitness of the immigrant strain in the range of high fitness values and (ii) the preferential emergence of only “lucky”, “fast-emerging” realizations in the range of low fitness values, lead to emergence time reaching a maximum in the stochastic regime.

Yield losses derived from the stochastic regime are lower than yield losses derived from the deterministic approach (Figs. 4e, f). This difference can be seen as the consequence of differences observed between these regimes both in terms of the probability of emergence (Figs. 4a, b) and the time to emergence (Figs 4c, d): above the deterministic threshold, there are cases where disease does not emerge, or where time to emergence is delayed in the stochastic approach, and therefore yield losses are not as high as in the deterministic approach.

### 4.3 Further questions to address on disease emergence

Our analyses provide a series of elements to better understand how disease emerges. The model structure allows addressing other important questions on disease emergence, such as:

- the effect of primary infection patterns on emergence: in the analyses we conducted, we considered only one type of primary infection, as a single immigration event occurring at a single point in time. The model allows the consideration of other patterns, including varying size of immigrant inoculum, or repeating inflows of immigrant strains over several cropping seasons (instead of during only one cropping season).
- the immigration rate simply considers the entry into the system of a new strain, with no specific hypothesis attached to the origin of this strain. The model also allows consideration of other potential sources of new strains, including recombinants or mutants, which could originate from inside or outside of the zone of emergence.
- the model can also include adaptation of the pathogen population, for example by varying *RRg* and *RRD* over time, or draw new parameters at the start of each cropping cycle according to the parameter values of the preceding cropping cycle (Figure 1, path 3).
- the effect of variation of *RRg* within the growing season can be analysed in order to mimic the effect of weather (e.g. warmer winters or drier summers) on epidemics and emergence (Figure 1, Path 2).
- the analyses were conducted with a relatively limited number of cropping cycles of simulation. This was appropriate because a large amount of inoculum was used in the simulations. When considering a low rate of immigration within a stochastic regime, much longer time frames may be needed to detect emergence.
- the effect of climate change on disease emergence can be addressed with this model by incorporating a directional change in the mean and/or the standard deviation of some fitness parameters (e.g., *RRg* and *RRD*) over successive cropping cycles.

Plant disease emergence is a complex phenomenon, with many system- and context-specific variants. This work addresses the phenomenon in a simplified manner in order to derive some of its main features. While this work needs to be continued, we hope that the present analysis provides a useful step towards implementing more effective policies to prevent or delay plant disease emergence.

## DATA AVAILABILTY STATEMENT

The data that support the findings of this study are available from the corresponding author upon reasonable request.

## Appendix A

Assuming that the dynamics of the two pathogen strains are independent of each other, the simulation model described in Sec. 2.2 [Eqs. (1)-(4)] can be summarised in a single equation:

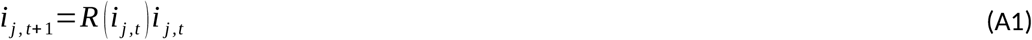

Where

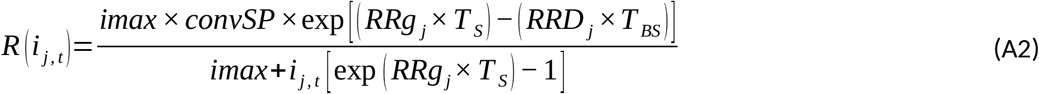

Here, *i*_j,t_ is the injury caused by the pathogen strain *j* (*j*=1 for the local strain, and *j*=2 for the immigrant strain) at the very beginning of the growing season *t*, where *t* is the index that runs through successive cropping cycles (i.e., *t* = 1,2,3…). *T*_S_ is the duration of the growing season (= *CGP*) and *T*_BS_ is the duration between two successive growing seasons. Eq. (A1) is a map that relates the injury at the beginning of growing season *t*+1, *i*_j,t+1_, to the injury at the beginning of the previous growing season *t*, *i*_j,t_, representing a nonlinear generalisation of the classical geometric growth model. The map Eq. (A1) has two fixed points:

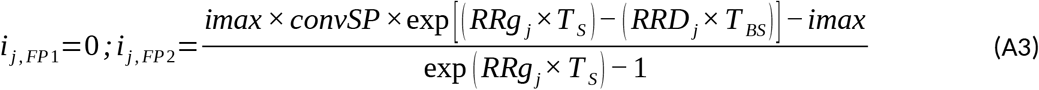

The fixed points determine the long-term outcomes of the dynamics: eventually the strain *j* either dies out (FP1) or reaches the stable equilibrium (FP2). The equilibrium occurs due to a balance between pathogen reproduction during the growing season and its decay between growing seasons: the number of newly produced pathogen individuals during the growing season compensates the number of individuals lost during the preceding between-growing season phase. Which of the two fixed points is achieved in the long run, is determined by the growth rate of the map Eq. (A1) linearised in the vicinity of FP1:

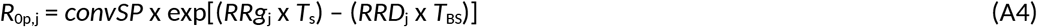

where *R*_0*p,j*_ quantifies the reproductive fitness of the strain *j* and corresponds here to the polyetic basic reproductive number. Usually, the basic reproductive number is defined as the number of host individuals that are infected by a single infected host introduced into a fully susceptible host population (e.g., Zadoks and Schein, 1979; Anderson and May, 1986; Campbell and Madden, 1990). Adapted to our context, the biological meaning of *R*_0p,j_ is the number of units of crop injury appearing at the beginning of growing season *t* + 1 following the introduction of a unit host injury at the beginning of the previous growing season *t.* If each of the two strains is viable when present alone, i.e., *R*_0p,j_ > 1, the strain that has a higher basic reproductive number eventually outcompetes the other strain. Consequently, the immigrant strain emerges if it has a higher polyetic basic reproductive number, i.e., *R*_0p,2_ > *R*_0p,1_. The emergence threshold is given by the equality of the two polyetic basic reproductive numbers: *R*_0p,2_ = *R*_0p,1_. We solve this equation with respect to *RRg*_2_, using Eq. (A4), to obtain the threshold value of *RRg*_2_ above which the emergence takes place:

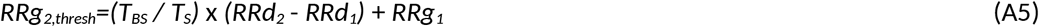

Similarly, the emergence threshold can be expressed in terms of *RRD*_*2*_:

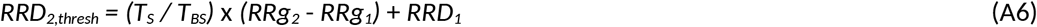

Here, the immigrant strain emerges when its relative decay rate is below the threshold, i.e., *RRD*_2_ < *RRD*_2,thresh_. When only one pathogen strain is present, *R*_0p,j_ given by Eq. (A4) determines without any approximation which of the two fixed points in Eq. (A3) will be achieved according to the map in Eq. (A1). However, when both pathogen strains are present, Eq. (A4) and Eqs. (A5), (A6) derived from it, give only approximate expressions for emergence thresholds, under the assumption that the saturation effects of the logistic growth are negligible. Nevertheless, *i*_j,FP2_ in Eq. (A3) provides an exact expression for the final, equilibrium level of injury due to the pathogen strain that wins the competition.

---

**Supplementary Figure 1.**
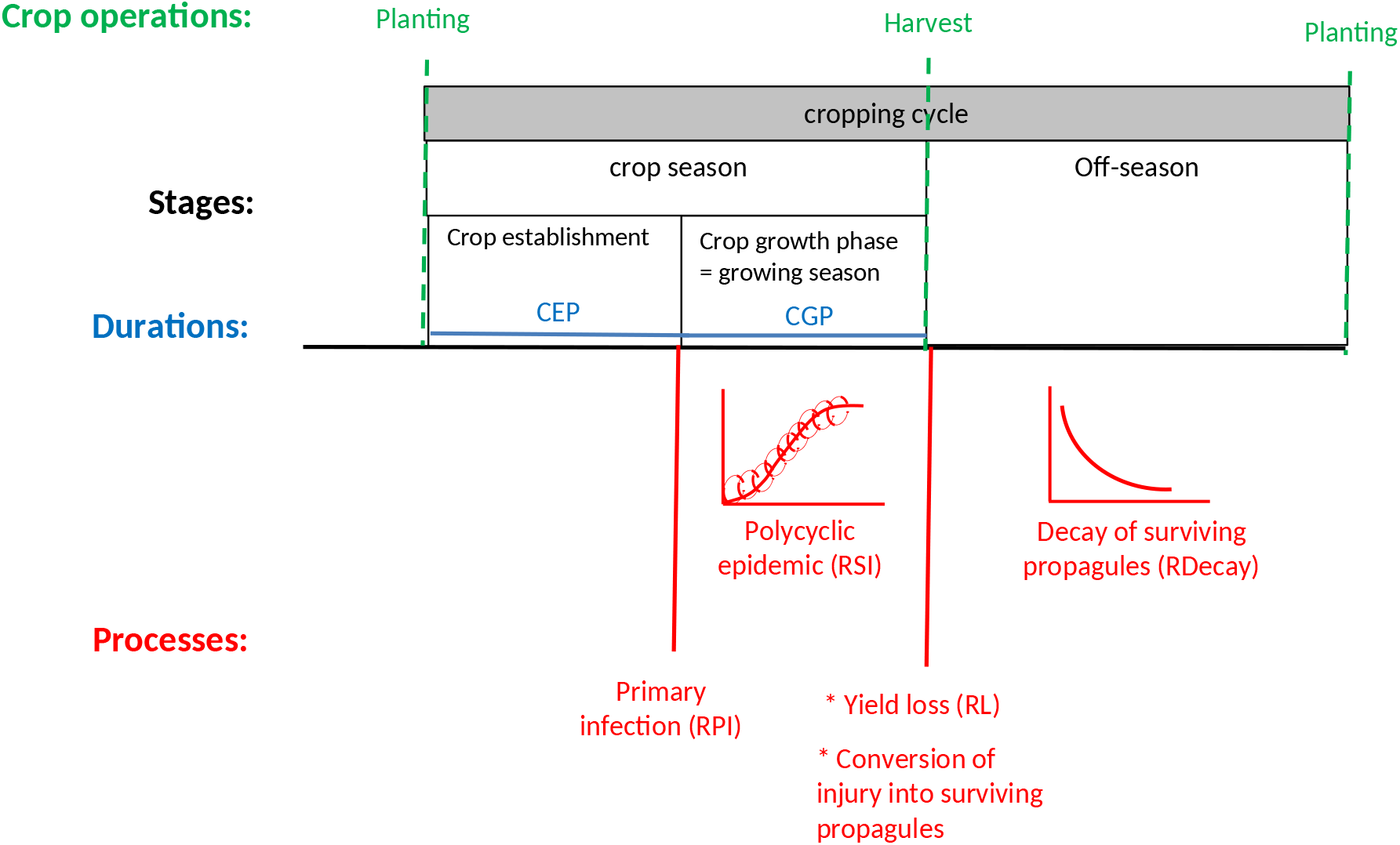
Main stages and processes considered in the polyetic model at each cropping cycle. CEP: duration of the Crop Establishment Phase; CGP: duration of Crop Growth phase.

